# Predicting RNA splicing from DNA sequence using Pangolin

**DOI:** 10.1101/2021.07.06.451243

**Authors:** Tony Zeng, Yang I Li

## Abstract

Recent progress in deep learning approaches have greatly improved the prediction of RNA splicing from DNA sequence. Here, we present Pangolin, a deep learning model to predict splice site strength in multiple tissues that has been trained on RNA splicing and sequence data from four species. Pangolin outperforms state of the art methods for predicting RNA splicing on a variety of prediction tasks. We use Pangolin to study the impact of genetic variants on RNA splicing, including lineage-specific variants and rare variants of uncertain significance. Pangolin predicts loss-of-function mutations with high accuracy and recall, particularly for mutations that are not missense or nonsense (AUPRC = 0.93), demonstrating remarkable potential for identifying pathogenic variants.

---

RNA splicing is an intricate gene regulatory mechanism that removes introns from pre-mRNAs. How the cell chooses which splice sites are used during RNA splicing remains unclear, but it is well known that DNA sequence is a key determinant of splice site usage (Senapathy et al., 1990; Blencowe, 2000; Wang et al., 2006). Many studies have now shown that both rare and common genetic variants contribute to human disease by disrupting RNA splicing (Li et al., 2016; Aguet et al., 2020). Thus, predicting RNA splicing from DNA sequences can greatly aid the identification and interpretation of disease-causing mutations.

Due to the complexity of the sequence determinants of RNA splicing, end-to-end deep neural networks are well-suited to learn features directly from DNA sequence to predict splicing outcomes of interest. Examples of state-of-the-art deep neural networks for predicting RNA splicing from DNA sequence include MMSplice (Cheng et al., 2019) and SpliceAI (Jaganathan et al., 2019), both of which have been used successfully to predict pathogenic mutations that impact splicing. Nevertheless, both methods have limitations. For example, MMSplice predicts usage of cassette exons rather than that of splice sites, and thus is likely to overlook disease-causing mutations that disrupt complex splicing patterns. Furthermore, neither MMSplice nor SpliceAI predicts tissue-specific splicing (though a newer version of MMSplice, MTSplice (Cheng et al., 2021) can predict tissue-specific splicing). Several methods that do not use deep learning have also been successfully used to study sequence determinants of splicing, including HAL (Rosenberg et al., 2015) and MaxEntScan (Yeo and Burge, 2004), which use an additive linear model and a maximum entropy model respectively. However, deep learning based approaches have been shown to outperform these methods in a variety of prediction tasks (Cheng et al., 2019; Jaganathan et al., 2019).

To model splicing in a quantitative, tissue-specific manner, we developed Pangolin, a deep neural network that predicts splicing in four tissues – heart, liver, brain, and testis – which represent major mammalian organs (Figure 1a). This is an improvement over SpliceAI and MMSplice, which produce the same prediction for any tissue. In addition, Pangolin can predict the usage of a splice site in addition to the probability that it is spliced (Figure 1a). This is an improvement over SpliceAI, which merely reports predictions for whether a dinucleotide is a splice site or not (Jaganathan et al., 2019) (Supplementary Note 1). Pangolin’s model architecture consists primarily of 16 stacked residual blocks with skip connections, each of which contains batch normalization, ReLU activation, and convolutional layers (Methods). Pangolin’s architecture resembles that used in SpliceAI, which allows modeling of features from up to 5,000 base pairs upstream and downstream each target splice site. An important difference is the addition of multiple outputs to the final neural network layer of Pangolin, allowing prediction of splice site usage across different tissues.

**Figure 1:**
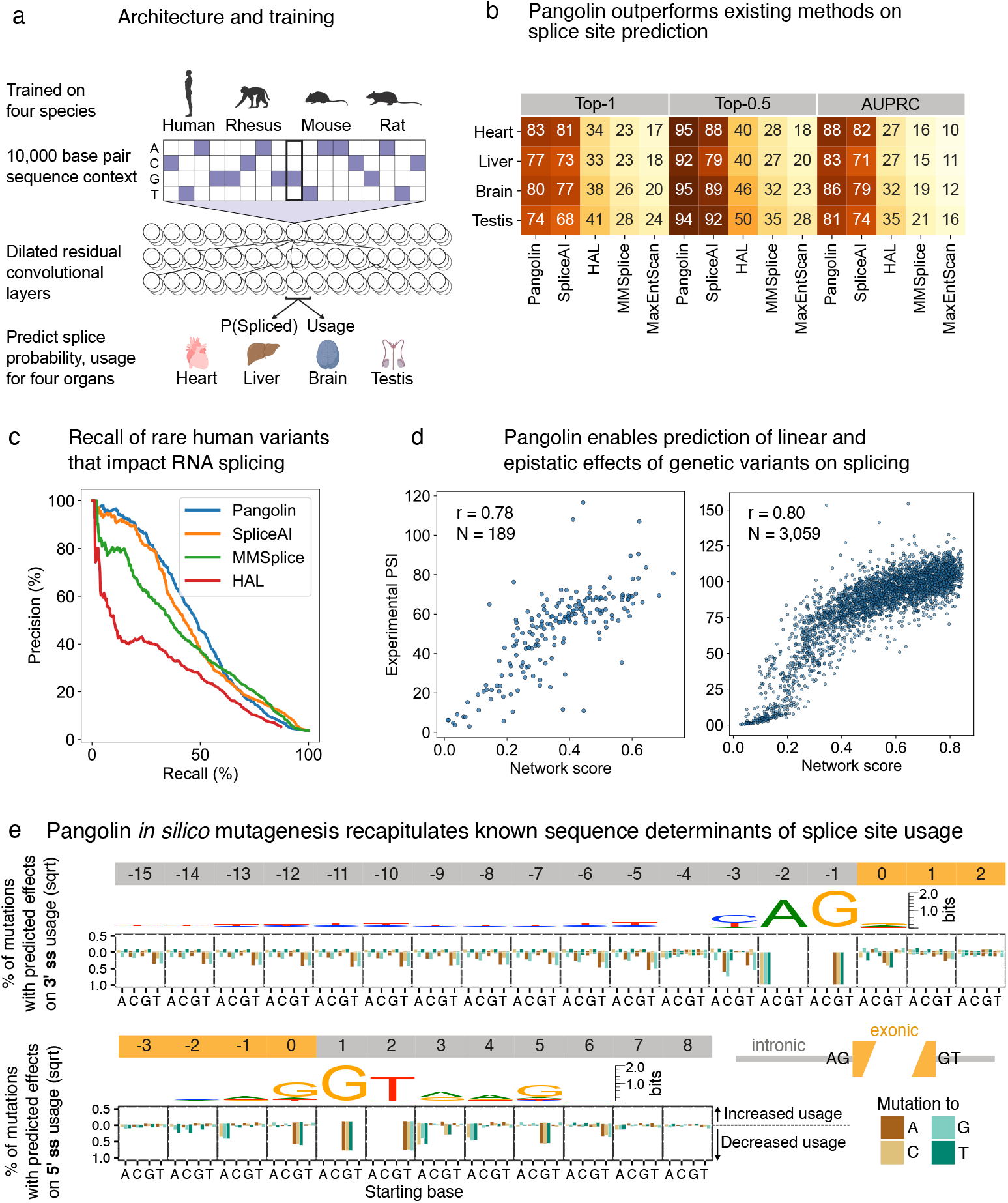
Overview of Pangolin and evaluation. (a) Schematic and architecture of Pangolin (b) Heatmap summarizing the performance of Pangolin, SpliceAI, HAL, MMSplice, and MaxEntScan with respect to three metrics including top-1 accuracy. (c) Precision-recall curves representing the precision and recall from multiple methods for the prediction of splice-disrupting variants as identified in Cheung et al., 2019 (1,050 splice-disrupting variants out of 27,733 total). (d) Scatter plots showing measured versus predicted effects of single genetic variants (left) or a combination of genetic variants (right) on RNA splicing. Measured effects of single genetic variants and combinations of variants were obtained from Julien et al., 2016 and Baeza-Centurion et al., 2019 respectively. (e) *In silico* mutagenesis of 6,416 exons from human chromosomes 7 and 8. Barplots show for each base the percent of mutations (square root) predicted to increase or decrease usage by at least 0.2.

To train Pangolin, we used data from four species—human, rhesus macaque, rat, and mouse—as we reasoned that prediction of splice site usages in multiple tissues may require a larger training set compared to that used in SpliceAI and MMSplice, which were trained using human sequences only. Using sequences from multiple species has been shown to substantially improve prediction in some applications (Kelley, 2020), but has not yet been tested, to the best of our knowledge, in the context of RNA splicing (Supplementary Note 2). Specifically, we processed sequence and RNA splicing measurements from the four aforementioned species (Figure 1a, Supplementary Table 1), by using SpliSER (Dent et al., 2020) to quantify the usage of all splice sites after mapping RNA-seq data from heart, liver, brain, and testis from up to 8 samples per species per tissue (Methods). Next, we split genes into a training set and a test set. The training set consists of genomic positions from genes on human chromosomes 2, 4, 6, 8, 10-22, X, and Y, including splice and non-splice sites, while the test set consists of positions from genes on human chromosomes 1, 3, 5, 7, and 9. To add training sequences from rhesus macaque, rat, and mouse, we used genes that do not show orthology or paralogy to genes from the test human chromosomes (Methods). This ensures that our model does not learn human patterns of RNA splicing from orthologous sequences.

We evaluated the performance of Pangolin on predicting splice sites alongside popular methods including MaxEntScan (Yeo and Burge, 2004), SpliceAI (Jaganathan et al., 2019), MMSplice (Cheng et al., 2019), and HAL (Rosenberg et al., 2015). We first compared all methods in terms of their splice site predictions on test chromosomes using the top-1 and top-0.5 metric (Methods). The top-1 (resp. top-0.5) metric measures the fraction of correctly predicted splice sites at the score threshold where the number of predictions equals the total number (resp. half the number) of labeled splice sites. We also used the area under the precision-recall curve (AUPRC) to assess performance on predicting splice site locations. Across all tissues tested, Pangolin achieved an average top-1 accuracy of 79% and AUPRC of 0.85—an improvement over SpliceAI, which achieved an average top-1 accuracy of 75% and AUPRC of 0.77. Both Pangolin and SpliceAI outperformed the other three tested methods—MMSplice, HAL, and MaxEntScan—which achieved average top-1 accuracies and AUPRC lower than 37% and 0.30 respectively (Figure 1b). When considering only the top half most confident splice site predictions, Pangolin also outperformed SpliceAI (94% vs 87% top-0.5 accuracy) and other methods (Figure 1b).

We next evaluated the methods in terms of their ability to predict the effects of rare variants on RNA splicing. We used data generated from a Sort-seq assay, MFASS, that tested the effects of 27,733 exonic and intronic variants from the Exome Aggregation Consortium (ExAC) on exon usage using minigene reporters (Cheung et al., 2019), most of which are extremely rare in the human population. About 3.8% of tested variants were found to strongly affect exon usage (Δinclusion index ≤ −0.5) and were therefore defined as splice-disrupting variants (SDVs) (Cheung et al., 2019). We tested the ability of Pangolin, SpliceAI, MMSplice, and HAL to distinguish SDVs from other variants. Pangolin achieved an AUPRC of 0.49, outperforming all other methods (Figure 1c, Methods), while SpliceAI scored the second best AUPRC (0.47). Most of the improvement in AUPRC came from higher precision at low and moderate rates of recall (precision of 87%, 81%, and 62% at 25% recall; and 46%, 39%, and 38% at 50% recall for Pangolin, SpliceAI, and MMSplice respectively). These results indicate that highly confident predictions (>80% precision) from Pangolin capture a substantial fraction (>25%) of rare variants that disrupt RNA splicing.

Pairwise and higher-order epistatic interactions between variants can frequently result in splicing outcomes that differ from those caused by single variants. To evaluate our ability to predict the effects of multiple mutations, we used Pangolin to predict percent-spliced-in (PSI) of exon 6 of the *FAS* gene with different single nucleotide substitutions (n = 189) or with a combination of multiple substitutions (n = 3,059) (Methods). We then compared our predictions to the experimental PSIs determined using minigene reporter assays (Julien et al., 2016, Baeza-Centurion et al., 2019). Spearman correlation between predicted and experimental PSI was high for single substitutions (*r*_single_ = 0.78) and even higher for combinations of substitutions (*r*_multiple_ = 0.80) (Figure 1d). By contrast, using a linear model of the individual variants’ PSIs (Baeza-Centurion et al., 2019) to predict the PSIs for combinations of variants resulted in a Spearman correlation of 0.48 (Supplementary Fig. 1, Methods). These results demonstrate Pangolin’s ability to account for epistatic effects when making predictions.

We also tested Pangolin’s performance on predicting tissue-specific splice site usage. To do this, we calculated the difference between estimated splice site usage in each tissue and the mean usages across tissues for each site in genes on the test chromosomes, then compared these observed differences to Pangolin’s predicted differences. We limited this analysis to sites whose usage in at least one tissue differed from the mean by > 0.2. Across tissues, the Spearman’s *r* coefficients between the observed and predicted tissue-specific splice site usages ranged from 0.35 to 0.50 (median of 0.43) (Supplementary Fig. 2). Thus, Pangolin is able to capture tissue-specific splicing effects to some extent. Although these correlations are low, we note that these are comparable to or higher than those of MTSplice (Cheng et al., 2021), which produced predictions for differential exon inclusion with Spearman’s *r* coefficients that ranged from 0.09 to 0.40 (median of 0.22) (Supplementary Note 3). We thus conclude that predicting differential splicing across tissues from sequence alone is possible but remains a considerable challenge.

Next, we applied Pangolin to a variety of prediction tasks as a demonstration of multiple potential use cases. As a first use case, we performed an *in silico* mutagenesis on human exons to visualize the effects of mutations on splice site usage (Methods). We predicted the effects on splice site usage of all possible mutations near the 3’ and 5’ splice sites for several thousand exons, and asked about the type and fraction of base changes that increase or decrease predicted usage by at least 0.2 (Methods). The mutational patterns predicted to impact splicing are highly consistent with known sequence motifs near splice sites (Figure 1e). For example, nearly all mutations away from the AG acceptor and GT donor dinucleotides are predicted to decrease usage of the 3’ and 5’ splice sites respectively. In addition, Pangolin predicts that for a large fraction of 3’ splice sites, upstream T to G or T to A mutations—and to a lesser extent T to C mutations—substantially reduce 3’ splice site strength. Conversely, upstream mutations to a T—and to a lesser extent mutations to a C—increase 3’ splice site strength. These predictions reveal the importance of polypyrimidine tract strength for the splicing of a substantial fraction of exons (Coolidge et al., 1997). Overall, we found that many fewer mutations are predicted to increase splice site usage than decrease it (Figure 1e). This finding suggests that most (but not all) exons harbor sequences that allow for near-optimal splicing accuracy. For example, we found that at the −3 position relative to the 3’ splice site, mutations away from C and T cause strong decreases in splice site usage (for another example at position −1 relative to the 5’ splice site, see Supplementary Note 4). This is consistent with a preference of the U2AF1 splicing factor to bind 3’ splice sites with C or T at the −3 position (Yoshida et al., 2020; Ilagan et al., 2015). Interestingly, a mutation to C or T at the −3 position often does not conversely increase usage, suggesting that 3’ splice sites with an A or G at the −3 position may be less reliant on U2AF1 binding for splice site recognition. These examples suggest that there exist prevalent mutational effects which depend on sequence context and cannot be captured by standard motifs or position weight matrices.

As a second use case, we used Pangolin to aid in the prediction of common genetic variants that impact RNA splicing. We asked Pangolin to identify single-nucleotide polymorphisms (SNPs) that impact intron excision at the top 500 most significant splicing quantitative trait loci (sQTL) identified from the DGN consortium using Leafcutter (Methods). We restricted analysis to sQTLs for which the lead sQTL was at most 1kb away from a splice site, and used Pangolin as well as SpliceAI to predict the effects on intron splice site usage for all SNPs within 1kb of splice sites (Methods). We reasoned that the *p*-value of the causal sQTL SNP should be generally of similar magnitude to that of the lead sQTL SNP. Thus, we compared the sQTL *p*-value of the SNP predicted to have the largest effect on splicing to the lead sQTL *p*-value. We found that SNPs predicted to be causal by Pangolin have sQTL *p*-values that are smaller than those predicted by SpliceAI, which in turn are much smaller than those of randomly chosen SNPs (Figure 2a).

**Figure 2:**
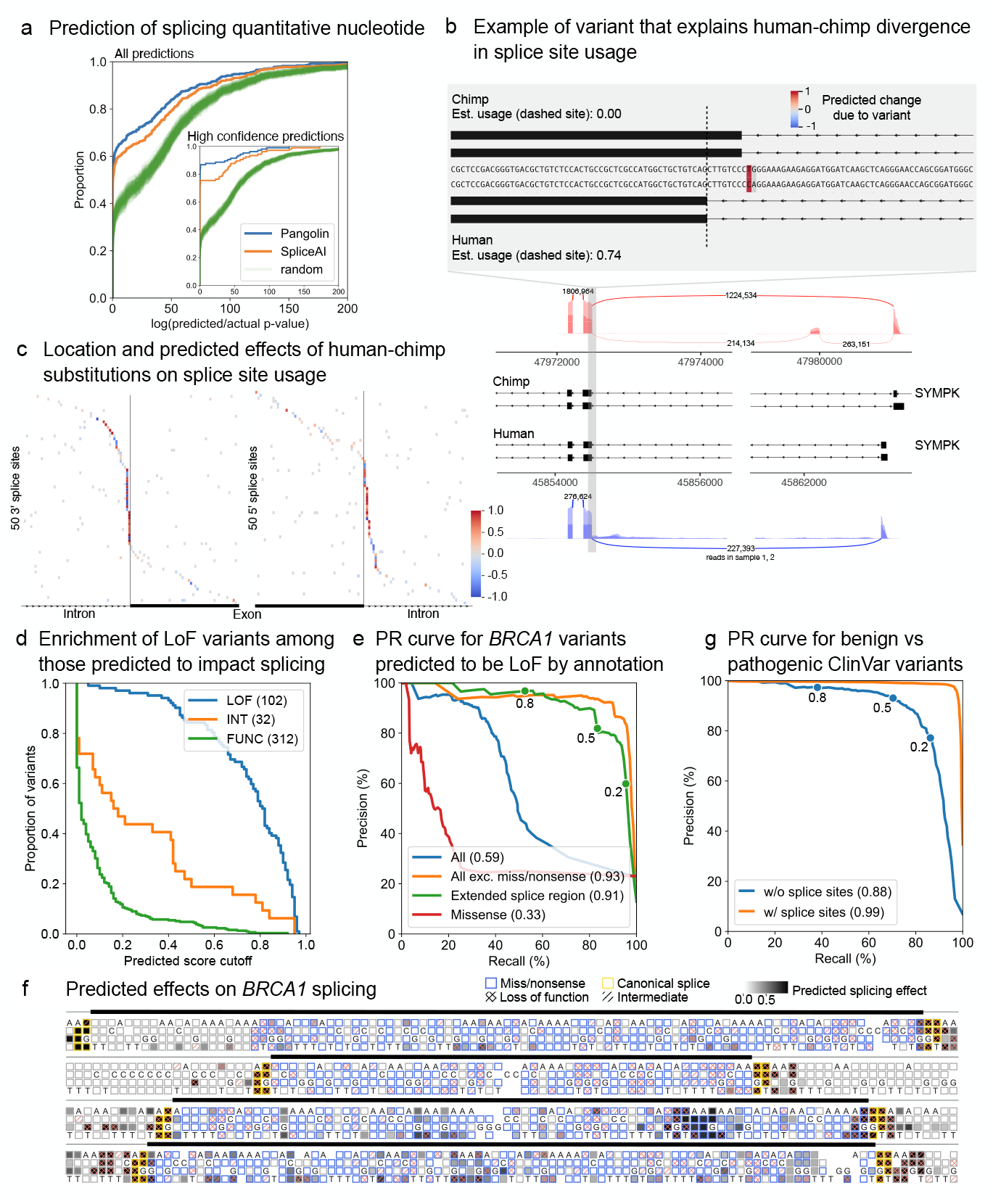
Application of Pangolin to a variety of prediction tasks. (a) Cumulative density plot of the log_10_ sQTL *p*-value fold difference between the SNP predicted to affect splicing and that of the lead sQTL SNP for the top 500 sQTLs identified in DGN (All predictions), or for the 106 predictions with the largest predicted effects (inset). (b) Example of a splice site that shows a large inter-species difference in usage. A single nucleotide difference between chimp (T) and human (C) is predicted to strongly decrease/increase usage of a chimp/human splice site (dashed vertical line indicates the human site). The T/C difference likely disrupts/creates a 3’ canonical splice site in chimp/human. (c) Locations and effects of SNVs ±50bp from a splice site predicted to underlie inter-species differences in splice site usage for 71 3’ and 74 5’ sites. A large fraction—but not all—of splice-altering variants are located near the canonical splice sites. (d) Survival function plots of *BRCA1* variants in splice regions as a function of their predicted effects on splicing. The variants are separated by their classification as loss-of-function (LOF, blue), intermediate effect (INT, orange), or functional (FUNC, green). We observe a huge enrichment of LOF variants among variants with large predicted splicing effects. (e) Precision-recall curves for different variant types representing the precision and recall for distinguishing LOF variants from functional variants. Pangolin achieves a remarkable AUPRC for variants in extended splice regions (note that this excludes canonical splice variants). See Supplementary Fig. 5 for variants from additional annotation bins. Precision and recall at three Pangolin score cutoffs are indicated. (f) Predicted splicing effects of mutations in or flanking 4 *BRCA1* exons from Findlay et al., 2018. Mutations identified to be LOF or to have intermediate phenotypes, as well as missense, nonsense, and canonical splice site mutations are annotated. See Supplementary Fig. 6 for all 13 exons with predictions. (g) Precision-recall curves representing the precision and recall for distinguishing variants annotated as pathogenic from variants annotated as benign in ClinVar. The blue (orange) line represents the PRC for variants excluding (including) variants in canonical splice sites. Missense and nonsense variants are excluded. Precision and recall at three Pangolin score cutoffs are indicated.

In particular, for 57% of sQTLs tested, Pangolin predicted a causal SNP with a *p*-value within one order of magnitude of the lead sQTL’s *p*-value (53% for SpliceAI, and 31% for random SNPs). Notably, this percentage increased to 87% for the 106 loci with the most confident Pangolin predictions (cutoff = 0.106, Methods) (Figure 2a, inset). These results suggest that SpliceAI and Pangolin are able to identify putatively causal SNPs for a substantial fraction of sQTLs, with Pangolin again outperforming SpliceAI.

As a third use case, we used Pangolin to study the genetic basis of inter-species variation in splice site usage—specifically, variation between human and chimp (note that Pangolin was not trained on chimp data). To this end, we first identified splice sites that are differentially used between human and chimp in brain tissue, focusing on splice sites with a large (≥ 0.5) difference in usage (Methods, Supplementary Table 2). We then asked Pangolin to predict these differences in usage, and found that a cutoff of 0.14 for the predicted differences resulted in a false sign rate of about 5% (Supplementary Note 5, Supplementary Fig. 3). Using this cutoff, we were able to identify 35% (550 out of 1,560) of the splice sites estimated to have differences in usage greater than 0.5, indicating that sequence divergence proximal (< 5kb) to the splice site contributes to a large fraction of human-chimp differences in RNA splicing. The remaining 65% of the splice sites with large differences in usage may be explained by the effects of mutations more than 5kb from the splice site or changes in the *trans-* cellular environment, or may represent false negatives that were not detected using Pangolin. Limiting further analysis to substitutions, we found that 47% (97 out of 206) of predicted differences can be explained by a single variant (Methods). A large fraction of these variants create or disrupt a canonical splice site (Figure 2b,c), as expected, but we also predict that many impact nearby sequences (Figure 2c). Thus, Pangolin can be used to pinpoint DNA variants responsible for the evolution of splice site usage.

As a last use case, we deployed Pangolin to predict the functional effects of mutations in known disease genes. We reasoned that mutations that are predicted to alter gene splicing are likely to impair function by decreasing functional isoform expression. Thus, accurate identification of mutations that impact splicing may aid in the functional interpretation of variants in disease genes, many of which would otherwise be of uncertain significance. To test this possibility, we first focused our analysis on *BRCA1* single-nucleotide variants whose functional impacts had previously been determined using saturation genome editing (3,893 variants in and around 13 exons of *BRCA1*, Findlay et al., 2018). We compared the effects of these variants on splicing as predicted using Pangolin to the functional impact of the variants as measured in Findlay et al., 2018. Strikingly, we found that variants determined to be loss-of-function (LOF) variants were highly enriched among variants predicted to impact RNA splicing (χ^2^ *p* = 5.6 × 10^−107^, Pangolin cutoff of 0.2), suggesting that impact on splicing is indeed informative for understanding functional impact. To identify the types of variants that drive this enrichment, we classified variants into four categories: non-synonymous, synonymous, splice region (8 intronic bases and 3 exonic bases flanking exon-intron boundaries, excluding non-synonymous and canonical splice variants), or intronic. We found that LOF variants were enriched among variants predicted to affect RNA splicing for all categories, including non-synonymous variants (χ^2^ *p* = 1.2 × 10^−23^, 2.1× enrichment, Supplementary Fig. 4), but particularly for variants in splice regions (χ^2^ *p* = 4.5 × 10^−25^, 3.0× enrichment, Figure 2d). Our findings indicate that although LOF variants in splice regions are the most likely to impact splicing, 5–10% of LOF missense variants may impact function through splicing effects rather than by altering protein sequence.

We next sought to directly evaluate Pangolin’s ability to predict LOF variants. This is a difficult task because only a small fraction (823 out of 3,893 tested *BRCA1* SNVs) of all possible variants are LOF, and this imbalance generally results in low precision for prediction tasks. Furthermore, many variants, including missense and nonsense variants, are expected to be LOF without affecting RNA splicing. Indeed, using Pangolin’s predicted splicing effects to distinguish LOF variants from functional variants results in a very low AUPRC for missense variants (AUPRC = 0.33). However, when excluding missense and nonsense variants, Pangolin achieves an AUPRC of 0.93 on the remaining 1,591 variants, and an AUPRC of 0.91 for the 861 variants in the extended splice region (±15bp from an exon-intron boundary excluding canonical splice variants, Figure 2e, Methods). Indeed, mutations predicted to have large impacts on RNA splicing of *BRCA1* appear to correlate particularly well with LOF status throughout the four shown exons (Figure 2f).

To generalize these findings, we applied Pangolin to variants in extended splice regions from the ClinVar database (Landrum et al., 2018), and found that Pangolin had a similar ability to distinguish the 833 SNVs labeled as pathogenic from the 10,999 SNVs labeled as benign (AUPRC = 0.88, Figure 2g, Methods), outperforming SpliceAI (AUPRC = 0.84, Supplementary Fig. 7). As expected, Pangolin’s performance improved when classifying splice region variants together with canonical splice site variants (AUPRC = 0.99, Figure 2g). Lastly, using Pangolin on 21,426 ClinVar SNVs labeled to be of unknown significance (VUS) in splice regions or canonical splice sites revealed that 5,173 VUS are likely to impact splicing and are thus likely pathogenic (Pangolin cutoff of 0.2, Supplementary Fig. 8 for *CHEK2* as an example, Supplementary Data 1 for Pangolin scores for all VUS). These results indicate that Pangolin can be used to identify non-missense and non-nonsense pathogenic variants with remarkable accuracy.

In conclusion, Pangolin outperforms contemporary methods for predicting RNA splicing from nearby DNA sequences, can be used for a variety of applications including pathogenic variant prediction, and is available freely online on GitHub (https://github.com/tkzeng/Pangolin).

## Methods

### Deep neural network architecture

Pangolin’s models are dilated convolutional neural networks with an architecture allowing features to be extracted from up 5,000 bases upstream and downstream each target site. It takes as input a one-hot encoded sequence of *N* bases, where *N* ≥ 10, 001, and predicts—for the middle *N* − 10, 000 bases—the probability that these sites are splice sites (probability outputs) and the usage of each site (usage outputs) in heart, liver, brain, and testis (see “Generating training and test sets” for details on the format of the input/output, and “Supplementary Note 1” for characterization of the two output types). Specifically, the neural networks consist of 16 stacked residual blocks, which are composed of batch normalization, ReLU activation, and convolutional layers; and skip connections, which add the model outputs before residual blocks 1, 5, 9, and 13 to the input of the penultimate layer. The convolutional layers of each block are dilated, allowing the receptive field width of the model to increase exponentially with the number of layers. The remaining components of the network are the first and penultimate layers, which are convolutional layers that transform the initial inputs and the residual block outputs respectively to dimensions appropriate for the subsequent layers; and the final activation functions that connect the penultimate layer to the output layer (softmax activations for the model’s probability predictions and sigmoid activations for the model’s usage predictions). In comparison to SpliceAI’s architecture, the final convolutional layers and activations are the primary points of difference—SpliceAI does not make probability/usage predictions for multiple tissues, but rather just makes one probability prediction (the probability that a site is a splice donor, splice acceptor, or not a splice site).

### Generating training and test sets

To identify splice sites and quantify their usages, we used RNA-seq data from four tissues—heart, liver, brain, and testis—across four species—human, rhesus macaque, mouse, and rat (Cardoso-Moreira et al., 2019). For heart, liver, and brain, we analyzed 8 samples for each species; for testis, we analyzed 8 samples for human, and 4 for rhesus macaque, mouse, and rat. Samples were chosen from developmental periods subsequent to all periods of large transcriptional changes (Cardoso-Moreira et al., 2019). RNA-seq reads were mapped to their respective genomes with annotations using STAR 2.7.5 (Dobin et al., 2013) in its multi-sample 2-pass mode (genomes and annotations: GRCh38 with GENCODE release 34 comprehensive annotations for human; Mmul 10 with ENSEMBL release 100 assembled chromosome annotations for rhesus macaque; GRCm38 with GENCODE release M25 annotations for mouse; and Rnor 6.0 with ENSEMBL release 101 assembled chromosome annotations for rat). Multimapped reads were assigned to a single location using MMR (Kahles et al., 2016).

To create training and test datasets for Pangolin for each tissue, we labeled every position within a gene body as spliced or not spliced, and quantified the usage of each splice site satisfying certain criteria. Specifically, we labeled all sites within gene bodies supported by 1 split read in at least 2 samples each as spliced, and we labeled all other sites as unspliced. To estimate usage levels of splice sites, we used SpliSER 1.3 (Dent et al., 2020) to calculate a Splice-Site Strength Estimate (SSE) for a subset of sites. We define the usage of a splice site as the proportion of transcripts of a gene that use a given splice site. SpliSER considers four types of reads to estimate usage for a target site: α and β_1_ reads are split and non-split reads respectively that map to the target site; and β_2−SIMPLE_ and β_2−CRYPTIC_ reads are split reads that provide direct and indirect evidence against usage respectively (Dent et al., 2020). Then, SSE is calculated as:

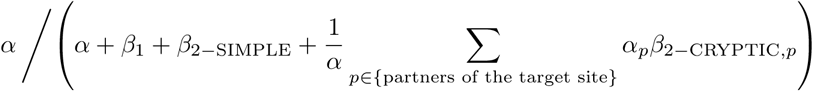

We estimated usage as SSE for all sites for which α + β_1_ + β_2−SIMPLE_ ≥ 5 in at least 2 samples each; sites that were labeled as spliced but did not meet these criteria were not assigned a usage estimate (so that usage predictions for these sites did not contribute to losses during training/testing), and all other sites were assigned a usage of 0.

We set aside genes from human chromosomes 1, 3, 5, 7, and 9 for the test set, and used for the training set all genes from human chromosomes 2, 4, 6, 8, 10-22, X, and Y that do not show paralogy to test set genes as well as all genes in rhesus macaque, mouse, and rat that do not show orthology to test set genes. To do this, we excluded all genes with either low or high “orthology confidence” to a test gene—determined using annotations from Ensembl BioMart (accessed 11/14/2020)—from the training set.

Next, we prepared the training set as follows so that the model could be trained in a sequence-to-sequence mode. For the model inputs, we extracted the sequence between the annotated left-most transcription start and right-most transcription end sites for each gene; zero-padded it so that the length was a multiple of 5,000 and that the start and end sites were each padded by 5,000 base pairs; and split the resulting sequence into blocks of 15,000 base pairs such that the *i*^th^ block contained positions 5,000(*i* − 1) − 5,000 to 5,000(*i* − 1) + 5,000. Each input sequence was then one-hot encoded: A, C, G, T/U, and—for unknown bases / padding—N were represented by [1, 0, 0, 0], [0, 1, 0, 0], [0, 0, 1, 0], [0, 0, 0, 1], and [0, 0, 0, 0] respectively. For the output labels, the middle 5,000 positions for each input sequence were each assigned a 12-dimensional vector: for tissues heart, liver, brain, and testis, positions 1-2, 4-5, 7-8, and 10-11 respectively were assigned one-hot encoded binary labels—[1, 0] for “is not splice site”, [0, 1] for “is a splice site”, and [0, 0] for unknown bases / padding—and positions 3, 6, 9, and 12 respectively were assigned the site’s estimated usage. For each target splice site, Pangolin’s output is also a 12-dimensional vector, containing predictions for the corresponding tissues and output types.

### Training Pangolin

During training, we randomly chose 10% of the 15,000 base pair blocks to determine an early-stopping point. We first trained the network using the AdamW optimizer and a warm restarts learning rate schedule with cycle lengths of 2 and 4 epochs (6 epochs total) (Loshchilov and Hutter, 2019). With this schedule, the learning rate was initialized to 5 × 10^−4^ at the start of each cycle and decayed to 0 with a cosine annealing by the end of each cycle. We computed losses for the network’s probability predictions using a categorical cross-entropy loss function, and losses for the network’s usage predictions using a binary cross-entropy loss function; the total loss for each input was computed as a sum over the losses for each prediction. The model was further trained on each tissue and label type (existence of splice site and splice site usage) separately, so that the loss for each input was computed using only one tissue and label type at a time (4 epochs for each tissue / label type; learning rate was initialized to 5 × 10^−4^ and decayed to 0 with a cosine annealing). In addition, the training procedure was run 5 times, resulting in 5 models per tissue / label type combination (40 total models). For all predictions of splice site probability or usage for a tissue, unless otherwise specified, we took the mean across the 5 models.

### Evaluation on held-out test set

For each tissue, we evaluated Pangolin’s ability to predict splice sites in genes from human chromosomes 1, 3, 5, 7, and 9, excluding genes with low expression levels (mean transcripts per million (TPM) across samples < 2.5 as determined using RSEM 1.3.3 (Li and Dewey, 2011)). We used the models trained to predict splice site probabilities for each tissue. We first computed for each tissue the average top-1 and top-0.5 accuracy over all genes from these chromosomes. In this setting, top-1 accuracy is the fraction of sites within the top *N* predicted splice sites that are labeled as splice sites, where *N* is the number of labeled splice sites in the test dataset. Similarly, top-0.5 accuracy is the same fraction but for the top ⌊*N/*2⌋ predicted splice sites. We also computed the area under the precision recall curve (AUPRC) for each tissue. For SpliceAI, we computed the probability that a site is spliced as the maximum of the 5’ and 3’ scores using the averages across the predictions of SpliceAI’s 5 trained models. For MMSplice (version 2.2.0), we applied the functions model.donorM.predict and model.acceptorM.predict (where model is a an instance of the MMSplice class) to input sequences 13bp into the intron and 5bp into the exon, and 50bp into the intron and 3bp into the exon—followed by a logit transformation—to predict the probability a site is a 5’ or 3’ site respectively. We then used the maximum of the 5’ and 3’ scores as the predicted probability that a site is spliced. For HAL, we used the function score seq pos (from Cell2015 N8 HAL Genome Predictions.ipynb in the GitHub repository https://github.com/Alex-Rosenberg/cell-2015) to obtain 5’ splice site scores, using 80bp into the intron and 80bp into the exon as input. Since HAL does not score 3’ splice sites, we labeled these sites as unspliced when evaluating HAL’s predictions. For MaxEntScan, we ran scripts score5.pl and score3.pl on input sequences 6bp into the intron and 3bp into the exon, and 20bp into the intron and 3bp into the exon—followed by the transformation 2^s^/ (2^s^ + 1) for each score *s*—to predict the probability a site is a 5’ or 3’ site respectively. We then used the maximum of the 5’ and 3’ scores as the predicted probability that a site is spliced. When running MMSplice, HAL, and MaxEntScan, we excluded inputs that contained ‘N’ bases (unknown bases or padding).

### Definition of maximum difference in scores

For some applications of Pangolin, we calculated the splice score of a variant as the maximum difference in scores between the reference and mutated sequence over a subset of the Pangolin’s models. We chose not to take the maximum over all 40 models because we found that introducing additional models beyond those in the subset did not affect performance much yet increased computational cost. Here, we define maximum difference in scores between two sequences. Let *P*_ref,tissue_ be the predicted probability that a splice site in the reference sequence context is spliced in a tissue, and *P*_alt,tissue_ be this probability for a splice site in the mutated sequence context. Let Δscores be the vector [*P*_alt,heart_ − *P*_ref,heart_, *P*_alt,liver_ − *P*_ref,liver_, *P*_alt,testis_ − *P*_ref,testis_]. Then, we define maximum difference in scores as the element in Δscores corresponding to max |Δscores|, i.e. Δscores[argmax|Δscores|].

### MFASS evaluation

Cheung et al., 2019 used a Sort-seq assay (MFASS) to quantify the effects of 27,733 exonic and intronic variants from the Exome Aggregation Consortium (ExAC) on exon recognition. These variants’ effects on splicing were assayed using minigene reporters each containing an exon and its surrounding intronic sequences, and variants with a Δinclusion index of ≤ −0.5 were classified as splice-disrupting variants. To score each variant using Pangolin, we computed—for the 5’ and 3’ sites—the maximum predicted difference in score between the mutated and reference sequences, and used the mean of the 5’ and 3’ differences as the score for the variant. Specifically, we used sequences ±5000bp of the 5’ and 3’ sites—obtained from the GRCh37 human reference assembly—as the reference sequence inputs to the model; and their mutated versions as the mutated sequence inputs. For SpliceAI, we scored each variant using the mean of the differences for the 5’ and 3’ sites between the reference and mutated sequence scores: 0.5 (score_5p,mut_ − score_5p,ref_ + score_3p,mut_ − score_3p,ref_). As before, the score for a sequence is the maximum of SpliceAI’s 5’ and 3’ scores. We computed precision and recall for Pangolin and SpliceAI using the precision recall curve function from scikit-learn, and AUPRC using the auc function. For MMSplice, we computed precision and recall using scripts from the MMSplice paper (Cheng et al., 2019, https://github.com/gagneurlab/MMSplice$_$paper), and for HAL, we used the precision and recall statistics from the MFASS paper (Cheung et al., 2019, https://github.com/KosuriLab/MFASS).

### *FAS* exon 6 evaluation

Julien et al., 2016 quantified the effects of all possible single mutations (189 total) in *FAS* exon 6 using a minigene reporter covering *FAS* exons 5-7. In a subsequent study, Baeza-Centurion et al., 2019 quantified the effects of several single, double, and higher-order combinations of 12 single mutations (3,072 total) in *FAS* exon 6. We used the first dataset to evaluate Pangolin’s performance on single mutations; and used sequences with > 1 mutation from the second dataset (3,059 out of 3,072) to evaluate Pangolin’s performance on multiple mutations. For the first dataset, we converted enrichment scores to PSI estimates by fitting an exponential calibration curve using 24 mutants with experimentally determined inclusion levels. For the second dataset, PSI estimates for each variant were provided in Baeza-Centurion et al., 2019. We scored each variant by computing max(*P*_heart_, *P*_liver_, *P*_testis_) for the 5’ and 3’ sites, where *P*_tissue_ is the predicted probability that a site is spliced in the specified tissue. We then used the mean of the scores for the 5’ and 3’ sites to predict the exon’s inclusion level for each variant. Reference sequences were obtained from the GRCh38 reference assembly. To understand the effects of epistatic interactions, Baeza-Centurion et al., 2019 developed a linear model with 12 parameters, one for each single mutation, to predict the PSIs of all exons in the library. We used this model to predict the PSIs for all exons with > 1 mutation, and calculated the Spearman’s *r* correlation coefficient between the predicted and observed PSIs.

### Tissue-specific splicing

To evaluate Pangolin’s ability to predict tissue-specific splicing differences, we considered a subset of sites in the test genes with higher confidence usage estimates: sites with at least 10 α+β_1_ +β_2_ reads per sample (see “Generating training and test sets” for definitions) for at least three samples and with standard deviations of < 0.1 for the usage estimates. Positions that did not meet these criteria but were labeled as splice sites (≥ 1 α read in ≥ 2 samples) were not considered; all other positions were set to have 0 usage. We also required that sites be expressed in all tissues (mean TPM across samples ≥ 2.5) and that for at least one tissue, the site’s usage differs from the mean usage across tissues by > 0.2. Then, for each tissue, we computed the Spearman’s *r* correlation coefficient between predicted and measured differences in usage for all sites in the test genes meeting the above criteria, where differences are taken between the usage in one tissue and the mean usage across all tissues.

### *In silico* mutagenesis

Using Pangolin, we evaluated for 5’ and 3’ sites the effects of all possible single base mutations 8bp into the intron and 4bp into the exon, and 15bp into the intron and 3bp into the exon respectively. We did this analysis for all exons from one representative transcript per gene (as determined by the presence of a MANE Select tag in the annotation file) for all protein-coding genes in human chromosomes 7 and 8, excluding the start of the first exon and end of the last exon of each transcript. We predicted the effect of each mutation on splice site usage by taking the mean across all tissues of the differences between usage predictions for the reference sequence and mutated sequence.

### Splicing QTLs evaluation

To evaluate Pangolin’s ability to predict the effects of common variants in their extant biological contexts, we used Pangolin to distinguish SNPs that are putatively causal for splicing differences, as determined using a splicing QTL (sQTL) analysis, from other nearby SNPs. We used a previously analyzed set of sQTLs generated using RNA-seq data from whole blood samples from 922 genotyped individuals in the Depression Genes and Networks (DGN) cohort (Mu et al., 2021). Briefly, Leafcutter was used to calculate intron excision ratios, which were used as the splicing phenotype, and sQTLs were called using fastQTL after accounting for principal component covariates. For each intron, we define a putatively causal SNP as the SNP with the most significant association (lowest *p*-value) with the splicing phenotype out of all tested SNPs. We considered QTLs whose causal SNPs were within 1,000bp of the 5’ or 3’ sites, and analyzed the 500 QTLs with the most significant causal SNPs. For each of these QTLs, we used Pangolin and SpliceAI to predict the splicing effects of the causal SNPs and all other SNPs within 1,000bp of the 5’ and 3’ sites that were considered during QTL calling. The predicted effect of each SNP was calculated as |Δ5’ score| or |Δ3’ score| if only the 5’ or 3’ sites respectively were within 1,000bp of the variant, and mean(|Δ5’ score|, |Δ3’ score|) if both sites were within 1,000bp of the variant. Here, Δ5’ score and Δ3’ score are the differences between the scores for the reference and mutated sequences (for Pangolin, we use the maximum differences in scores as defined previously).

Next, we generated empirical CDF (eCDF) plots for the ratio (predicted *p*-value)/(putative *p*-value), where predicted *p*-value for a given QTL is the *p*-value of the SNP with the largest predicted effect on splicing (out of all evaluated SNPs, as determined by Pangolin or SpliceAI), and putative *p*-value is the *p*-value of the putatively causal SNP. In such a plot, each point (*x, y*) on the graph represents the proportion of SNPs *y* for which the aforementioned ratio is less than *x*. As a baseline, we also generated eCDF plots for the ratio (random *p*-value)/(putative *p*-value), where random *p*-value for a given QTL is the *p*-value of a SNP uniformly drawn from the set of SNPs within 1000bp of the 5’/3’ sites; we generated 100 such plots. We considered a prediction to be high-confidence if the predicted effect on splicing was greater than cutoffs 0.106 for Pangolin and 0.1 for SpliceAI; these cutoffs were chosen so that an equal number of predictions passed the respective cutoffs for both Pangolin and SpliceAI.

### Splice site evolution

To predict variants responsible for differences in splice site usage between species, we analyzed RNA-seq data from human, chimpanzee, and rhesus macaque prefrontal cortex samples (Kanton et al., 2019). We analyzed two samples per species (one sample per individual). We combined RNA-seq reads from cortex sections for each individual and mapped reads to genome assemblies with annotations using STAR 2.7.5 (multi-sample 2-pass mode) (GRCh38 with GENCODE release 34 comprehensive annotations for human; Mmul 10 with ENSEMBL release 100 assembled chromosome annotations for rhesus macaque; and Pan tro 3.0 with ENSEMBL release 101 assembled chromosome annotations for chimpanzee). To convert coordinates between genomes, we used Liftoff (Shumate and Salzberg, 2021), a genome annotation lift-over tool, to map genomic features such as exons, transcripts, and genes from the human genome assembly to chimpanzee and rhesus macaque genome assemblies, and vice versa. We calculated usage for sites with at least 50 α + β_1_ + β_2_ reads in each sample (see “Generating training and test sets” for definitions) and with standard deviations of < 0.05 for the usage estimates. Positions that did not pass these criteria but had ≥ 1 α read in ≥ 1 sample were not considered; all other positions were set to have 0 usage. We also required that sites be in genes with mean TPM across samples ≥ 2.5. For comparisons of splice site usage between human and chimpanzee, we considered annotated human (chimpanzee) sites that mapped to the chimpanzee (human) genome with sequence identity ≥ 0.75 and coverage ≥ 0.75, as determined by Liftoff, and required the genes’ strands to be the same; sites that mapped to multiple locations, and locations that were mapped to multiple times were filtered out. We considered sites to be differentially spliced if |usage in human − usage in rhesus| ≥ 0.5. To score each differentially spliced site, we computed the maximum difference in splice scores between chimp and human; using these predictions, we calculated false sign rates (FSR) for a range of predicted score cutoffs (Supplementary Note 5).

To identify sites where a single mutation is sufficient to explain the difference in splice scores, we first limited analysis to sites predicted to be differentially used (5% FSR, score cutoff = 0.14) for which chimpanzee and human sequences showed at most 10% divergence in regions near the splice site (20 differences within 100bp upstream and downstream of the splice site). These nearby sequence differences are likely substitutions. Next, we kept sites where the predicted difference in usage is explained mostly by these nearby differences rather than by more distal ones. Specifically, let Ch denote the chimp sequence ±5000bp of the splice site and Hu denote the human sequence ±5000bp of the splice site but with the site’s local neighborhood (±100bp of the site) replaced by the chimp sequence (Supplementary Fig. 9). Also, let ChMutAll and HuMutAll be the Ch and Hu sequences but with the introduction of all human variants ±100bp of the splice site. We then kept sites where |score_Ch_ − score_Hu_| < δ and |score_ChMutAll_ − score_HuMutAll_| < δ, where δ is a small number defined as min (0.1, |score_ChMutAll_ − score_HuMutAll_|/ 5) (Supplementary Fig. 9). Next, we identified sites where a single nearby mutation is sufficient to explain the difference in splice scores regardless of the distal sequence context. Specifically, for each mutation ±100bp of the splice site, denoted as MutX, we check if |score_ChMutX_ − score_HuMutAll_| < δ and |score_HuMutX_ − score_HuMutAll_| < δ, where ChMutX and HuMutX are the Ch and Hu sequences respectively but with the introduction of MutX (Supplemental Fig. 9). We classify a site as having a single putatively causal mutation if only one mutation satisfies these inequalities. Examples of such sites are shown in Figure 2b and Figure 2c.

### *BRCA1* evaluation and ClinVar prediction

We applied Pangolin to 3,893 SNVs in 13 exons and nearby intronic regions of the *BRCA1* gene for which Findlay et al., 2018 performed saturation genome editing to determine their functional consequences (briefly, they performed editing in a cell line in which *BRCA1* is essential and calculated variant function scores using the variants’ depletion levels over time). For each variant, we used Pangolin to compute the largest decrease in splice score within 50 bases of each variant: min {min(Δscores_variant pos−50_), …, min(Δscores_variant pos+50_)}, where Δscores is as previously defined (Definition of maximum difference in scores). Input sequences were obtained from the GRCh37 reference assembly. Variants were classified as missense, nonsense, intronic, synonymous, splice region, or canonical splice variants using labels from Findlay et al., 2018. In particular, splice regions are sequences 3bp into the exon or 8bp into the intron that do not disrupt canonical splice sites or alter the amino acid sequence. Extended splice regions are defined similarly, but include sequences ±15bp of the exon-intron boundary. Variants were classified as loss of function (LOF), intermediate, or functional using the function score thresholds determined in Findlay et al., 2018; function scores were bimodally distributed, with only 6.4% of variants classified as intermediate. To compute precision-recall curves and AUPRC, we used Pangolin scores to classify variants as either LOF or functional.

We applied Pangolin to ClinVar variants downloaded from https://ftp.ncbi.nlm.nih.gov/pub/clinvar/vcf_GRCh38/clinvar.vcf.gz on 05/04/2021. Pangolin v0.0.1 was run on all variants using default settings: for all variants passing certain criteria, we computed the maximum increase and decrease in splice score within 50 bases of the variant using the GRCh38 genome assembly and GENCODE Release 34 basic gene annotations as inputs to the command-line program. These criteria were: the variant is a substitution or simple insertion/deletion (insertion/deletion where either the REF or ALT field is a single base); is contained in a gene body, as defined by the annotation file; is not within 5,000 bases of the chromosome ends; and is not a deletion larger than twice the input parameter -d. SpliceAI 1.3 was also run on the ClinVar VCF using default settings using the GRCh38 genome assembly and -A grch38 as inputs/parameters; SpliceAI scores variants satisfying similar criteria as those used by Pangolin. For further analyses, we considered variants in protein-coding genes that were either classified by ClinVar as Benign, Pathogenic, Likely Benign, Likely Pathogenic, or Uncertain significance; and required that each variant be in only one gene, not be a nonsense or missense variant as determined using the molecular consequence (MC) field (variants with no such field were not considered), and be within 15bp of an annotated splice site (this excludes the start of the first exon / end of the last exon of each transcript). We also only considered variants that had both Pangolin and SpliceAI scores. Variants that affect the canonical splice sites were identified using the MC field.

## Author’s contributions

Y.I.L. conceived of the project. T.Z. performed the analyses and implemented the software. T.Z. and Y.I.L. wrote the manuscript.

## Acknowledgements

We thank Benjamin Fair and Xuanyao Liu for comments on the manuscript.

## Availability of Data and Materials

RNA-seq data are available (ArrayExpress E-MTAB-6798 (mouse), E-MTAB-6811 (rat), E-MTAB-6813 (rhesus macaque), E-MTAB-6814 (human), E-MTAB-8231 (rhesus macaque, chimpanzee, human used in the splice site evolution analysis)), MFASS data are available (https://github.com/KosuriLab/MFASS), *FAS* exon 6 data are available (Supplementary Data 1 of Julien et al., 2016), DGN data are available (Mu et al., 2021), *BRCA1* data are available (Supplementary Table 1 of Findlay et al., 2018), ClinVar variants with Pangolin annotations are available (Supplementary Data 1), Pangolin is available on GitHub (https://github.com/tkzeng/Pangolin).

## References

Aguet, F., Anand, S., Ardlie, K. G., Gabriel, S., Getz, G. A., Graubert, A., Hadley, K., Handsaker, R. E., Huang, K. H., Kashin, S., et al., 2020. The GTEx Consortium atlas of genetic regulatory effects across human tissues. Science, 369(6509):1318–1330.

Baeza-Centurion, P., Miñana, B., Schmiedel, J. M., Valcárcel, J., and Lehner, B., 2019. Combinatorial Genetics Reveals a Scaling Law for the Effects of Mutations on Splicing. Cell, 176(3):549–563.

Blencowe, B. J., 2000. Exonic splicing enhancers: mechanism of action, diversity and role in human genetic diseases. Trends Biochem Sci, 25(3):106–110.

Cardoso-Moreira, M., Halbert, J., Valloton, D., Velten, B., Chen, C., Shao, Y., Liechti, A., Ascenção, K., Rummel, C., Ovchinnikova, S., et al., 2019. Gene expression across mammalian organ development. Nature, 571(7766):505–509.

Cheng, J., Çelik, M. H., Kundaje, A., and Gagneur, J., 2021. MTSplice predicts effects of genetic variants on tissue-specific splicing. Genome Biol, 22(1):94.

Cheng, J., Nguyen, T. Y. D., Cygan, K. J., Çelik, M. H., Fairbrother, W. G., Avsec, Ž., and Gagneur, J., 2019. MMSplice: modular modeling improves the predictions of genetic variant effects on splicing. Genome Biol, 20(1):48.

Cheung, R., Insigne, K. D., Yao, D., Burghard, C. P., Wang, J., Hsiao, Y. E., Jones, E. M., Goodman, D. B., Xiao, X., and Kosuri, S., et al., 2019. A Multiplexed Assay for Exon Recognition Reveals that an Unappreciated Fraction of Rare Genetic Variants Cause Large-Effect Splicing Disruptions. Mol Cell, 73(1):183–194.

Coolidge, C. J., Seely, R. J., and Patton, J. G., 1997. Functional analysis of the polypyrimidine tract in pre-mRNA splicing. Nucleic Acids Res, 25(4):888–896.

Dent, C., Singh, S., Mishra, S., Shamaya, N., Loo, K. P., Sarwade, R. D., Harrison, P., Sureshkumar, S., Powell, D., and Balasubramanian, S., et al., 2020. Splice-site strength estimation: A simple yet powerful approach to analyse rna splicing. bioRxiv,.

Dobin, A., Davis, C. A., Schlesinger, F., Drenkow, J., Zaleski, C., Jha, S., Batut, P., Chaisson, M., and Gingeras, T. R., 2013. Star: ultrafast universal rna-seq aligner. Bioinformatics, 29(1):15–21.

Findlay, G. M., Daza, R. M., Martin, B., Zhang, M. D., Leith, A. P., Gasperini, M., Janizek, J. D., Huang, X., Starita, L. M., and Shendure, J., et al., 2018. Accurate classification of BRCA1 variants with saturation genome editing. Nature, 562(7726):217–222.

Ilagan, J. O., Ramakrishnan, A., Hayes, B., Murphy, M. E., Zebari, A. S., Bradley, P., and Bradley, R. K., 2015. U2AF1 mutations alter splice site recognition in hematological malignancies. Genome Res, 25(1):14–26.

Jaganathan, K., Kyriazopoulou Panagiotopoulou, S., McRae, J. F., Darbandi, S. F., Knowles, D., Li, Y. I., Kosmicki, J. A., Arbelaez, J., Cui, W., Schwartz, G. B., et al., 2019. Predicting Splicing from Primary Sequence with Deep Learning. Cell, 176(3):535–548.

Julien, P., Miñana, B., Baeza-Centurion, P., Valcárcel, J., and Lehner, B., 2016. The complete local genotype–phenotype landscape for the alternative splicing of a human exon. Nature Communications, 7(1):11558. Number: 1 Publisher: Nature Publishing Group.

Kahles, A., Behr, J., and Rätsch, G., 2016. MMR: a tool for read multi-mapper resolution. Bioinformatics, 32(5):770–772.

Kanton, S., Boyle, M. J., He, Z., Santel, M., Weigert, A., Sanchís-Calleja, F., Guijarro, P., Sidow, L., Fleck, J. S., Han, D., et al., 2019. Organoid single-cell genomic atlas uncovers human-specific features of brain development. Nature, 574(7778):418–422. Bandiera abtest: a Cg type: Nature Research Journals Number: 7778 Primary atype: Research Publisher: Nature Publishing Group Subject term: Developmental neurogenesis;Evolutionary developmental biology;Stem-cell differentiation Subject term id: developmental-neurogenesis;evolutionary-developmental-biology;stem-cell-differentiation.

Kelley, D. R., 2020. Cross-species regulatory sequence activity prediction. PLoS Comput Biol, 16(7):e1008050.

Landrum, M. J., Lee, J. M., Benson, M., Brown, G. R., Chao, C., Chitipiralla, S., Gu, B., Hart, J., Hoffman, D., Jang, W., et al., 2018. ClinVar: improving access to variant interpretations and supporting evidence. Nucleic Acids Res, 46(D1):D1062–D1067.

Li, B. and Dewey, C. N., 2011. RSEM: accurate transcript quantification from RNA-Seq data with or without a reference genome. BMC Bioinformatics, 12(1):323.

Li, Y. I., Van De Geijn, B., Raj, A., Knowles, D. A., Petti, A. A., Golan, D., Gilad, Y., and Pritchard, J. K., 2016. Rna splicing is a primary link between genetic variation and disease. Science, 352(6285):600–604.

Loshchilov, I. and Hutter, F., 2019. Decoupled Weight Decay Regularization. arXiv:1711.05101 [cs, math],. arXiv: 1711.05101.

Mu, Z., Wei, W., Fair, B., Miao, J., Zhu, P., and Li, Y. I., 2021. The impact of cell type and context-dependent regulatory variants on human immune traits. Genome Biology, 22(1):122.

Rosenberg, A. B., Patwardhan, R. P., Shendure, J., and Seelig, G., 2015. Learning the sequence determinants of alternative splicing from millions of random sequences. Cell, 163(3):698–711.

Senapathy, P., Shapiro, M. B., and Harris, N. L., 1990. Splice junctions, branch point sites, and exons: sequence statistics, identification, and applications to genome project. Methods Enzymol, 183:252–278.

Shumate, A. and Salzberg, S. L., 2021. Liftoff: accurate mapping of gene annotations. Bioinformatics, (btaa1016).

Wang, Z., Xiao, X., Van Nostrand, E., and Burge, C. B., 2006. General and specific functions of exonic splicing silencers in splicing control. Mol Cell, 23(1):61–70.

Yeo, G. and Burge, C. B., 2004. Maximum entropy modeling of short sequence motifs with applications to RNA splicing signals. J Comput Biol, 11(2-3):377–394.

Yoshida, H., Park, S. Y., Sakashita, G., Nariai, Y., Kuwasako, K., Muto, Y., Urano, T., and Obayashi, E., 2020. Elucidation of the aberrant 3’ splice site selection by cancer-associated mutations on the U2AF1. Nat Commun, 11(1):4744.

