## Supplementary Fig. 1-11, Table 3 for "Predicting RNA splicing from DNA sequence using Pangolin"

### Supplementary Materials for
### Predicting RNA splicing from DNA sequence using Pangolin

#### Supplementary Notes

##### Supplementary Note 1: Binary versus continuous prediction output

SpliceAI outputs the probability that a dinucleotide is a splice site. We sought to test whether directly predicting splice site usage can improve prediction of splice site usage as compared to using the probability that a dinucleotide is a splice site as usage. To do this, we first placed sites into bins of usage [0, 0.1], (0.1, 0.5], (0.5, 0.9], and (0.9, 1]. For sites in adjacent bins (for example, sites with usage between (0.1, 0.5] or between (0.5, 0.9]), we used Pangolin's probability and usage outputs to predict which of the two bins the sites belonged to (Supplementary Fig. 10). Directly predicting splice site usage led to an improvement in AUPRC of 0.018 to 0.050 (median: 0.025). Violin plots show differences in the distributions of the probability and usage predictions for each bin (Supplementary Fig. 10). Depending on the application, we used either Pangolin's probability or usage predictions (Methods).

##### Supplementary Note 2: Training Pangolin using splicing data from multiple species improve prediction

In addition to training Pangolin as described in Methods (using human genes from chromosomes 2, 4, 6, 8, 10-22, X, and Y, and genes from rhesus macaque, mouse, and rat without paralogy or orthology to human genes on the test chromosomes), we also trained Pangolin on only human sequences. We trained two models separately for 13 epochs each (warm restarts learning rate schedule with cycle lengths of 2, 4, and 7 epochs). To evaluate differences in performance, we compared AUPRC for the task of classifying sites as spliced or unspliced, and found that the improvement in AUPRC from using all species compared to human only was small but significant (improvement in AUPRC of 0.013 to 0.025) for all tissues (Supplementary Table 3).

##### Supplementary Note 3: Pangolin versus MTSplice on predicting tissue-type-specific splicing

Cheng et al., 2021 developed a deep learning model, MTSplice, for predicting tissue-specific effects of variants on splicing. MTSplice consists of MMSplice (Cheng et al., 2019) and TSplice, a deep learning model that considers 100 bases into the exon and 300 bases into the neighboring introns to predict tissue-specific percent spliced-in (PSI). Cheng et al., 2021 evaluated TSplice by computing—for each tissue—the Spearman's $r$ correlation coefficient between the observed and predicted log odds ratios of tissue specific PSI for test set exons for which PSI deviated from the tissue-averaged

PSI by at least 0.2 in at least one tissue and for which the corresponding gene is expressed in at least 10 tissues (out of 51 total tissues). We evaluated Pangolin in an analogous fashion, but measured Spearman correlation for observed and predicted differences in splice site usage rather than differences in PSI (Methods). Due to differences in the phenotypes being predicted; the number of tissues evaluated (4 for Pangolin, 51 for MTSplice); the datasets used for evaluation; and other differences in implementation, Pangolin's correlations are not directly comparable to those of TSplice.

#### Supplementary Note 4: Mutations away from G at the -1 position of the 5' splice site cause strong decreases in 5' splice site usage

The G is complementary with U1 snRNA at the -1 position. Interestingly, the effect of mutating away from G does not reciprocate mutating to a G. We hypothesize that mutations on well spliced introns in general have more room to decrease their splicing efficiency than increase it. This observation also holds for the -3 position at the 3' splice site.

#### Supplementary Note 5: Predicting causal variants that explain inter-species divergence in splice site usage

To predict causal sequence differences underlying inter-species divergence in splicing, we used Pangolin to predict the effects of human-chimpanzee sequence differences near splice sites with large differences in usage between human and chimpanzee. We calculated a false sign rate (FSR) for different Pangolin score thresholds, which corresponds to the fraction of sites for which the predicted differences in splice site usage are of opposite directions from those of the observed differences. We found that at a fixed FSR, Pangolin was consistently able to predict the correct directions of effects (and thus the likely causal variants) for more splice sites in comparison to SpliceAI (Supplementary Fig. 11).

As validation, we sought to estimate the splice site usage of human-chimpanzee divergent sites in rhesus macaque. We reasoned that if the mutation predicted to explain human-chimp divergence in splicing occurred in the human (or chimp) lineage, then splice site usage in rhesus macaque would be closer to that in chimp (or human). To test this, we first obtained 46 differentially used splice sites (5% FSR, cutoff = 0.14) for which chimpanzee, human, and rhesus macaque sequences showed at most 10% divergence in regions near the splice site (20 differences within 100bp upstream and downstream of the splice site, pairwise comparisons between species). Out of these, we identified 17 sites where a single mutation sufficiently explains the predicted human-chimp difference in usage (Methods); for 16 of these sites, the rhesus macaque sequence at the location of the causal variant matched either the human or chimpanzee sequence. We found that for 14 of these 16 splice sites (88%), the predicted differences in splice site usage between human and chimpanzee were consistent with splice site usage measured in rhesus macaque. We considered a prediction to be consistent if the mutation occurred in the human lineage and the splice site's usage in rhesus is more similar to that in chimp than that in human, or the mutation occurred in the chimp lineage and the splice site's usage in rhesus is more similar to that in human than that in chimp.

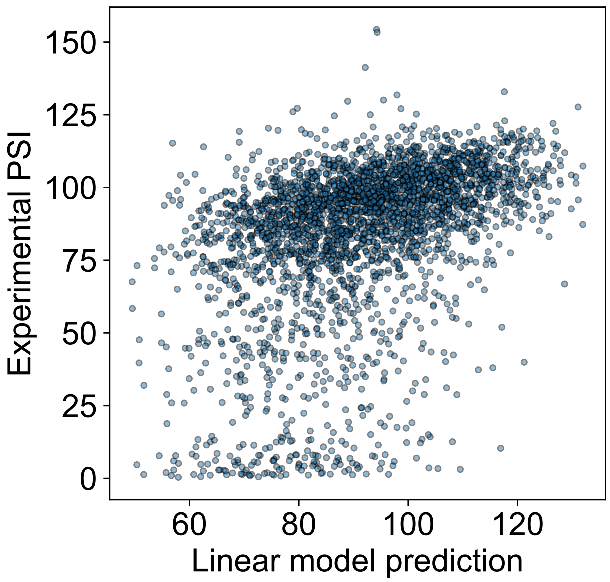


**Supplementary Figure 1. Prediction of epistatic effects on RNA splicing as a combination of single SNP effects**. Scatter plot showing measured (y-axis) versus predicted (x-axis) effects of combinations of genetic variants on RNA splicing. Measured effects of combinations of variants were obtained from Baeza-Centurion et al., 2019. Predictions were made using a linear model of the single variant PSIs (see Methods).

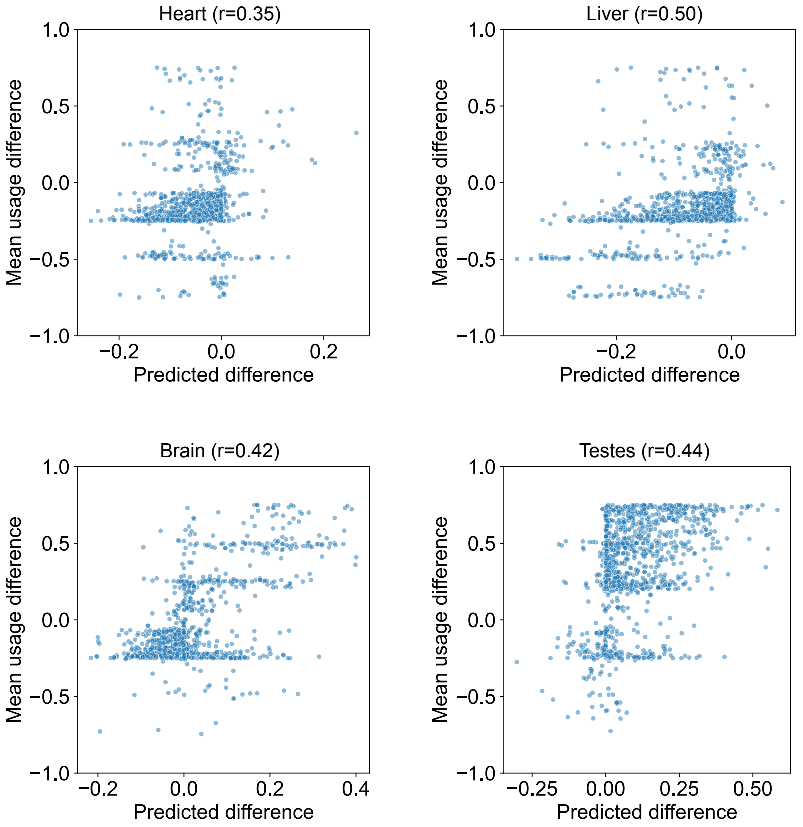


**Supplementary Figure 2. Prediction of tissue-specific splice site usage using Pangolin**. Scatter plots showing the mean empirical difference between splice site usage in the specified tissue and mean usage across all tissues (y-axis) versus the predicted difference as determined using Pangolin (x-axis). The prediction accuracies, although low, outperform or are comparable to those of MTSplice (Cheng et al., 2021). Our results suggest that predicting tissue-specific splicing remains challenging.

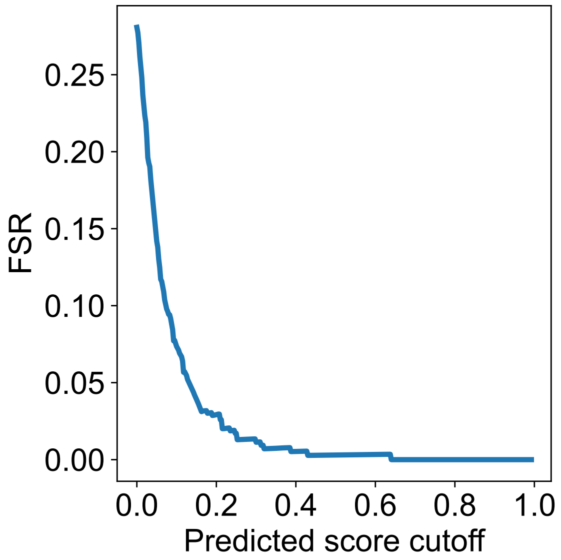


**Supplementary Figure 3. False sign rates of predicted causal variants underlying inter-species divergence in splice site usage**. False sign rates (FSR) at different cutoffs for predicted scores determined using Pangolin, calculated across 1,560 splice sites with large differences in usage ($\geq 0.5$) between human and chimpanzee. We observed a FSR of about 5% at a cutoff of 0.14.

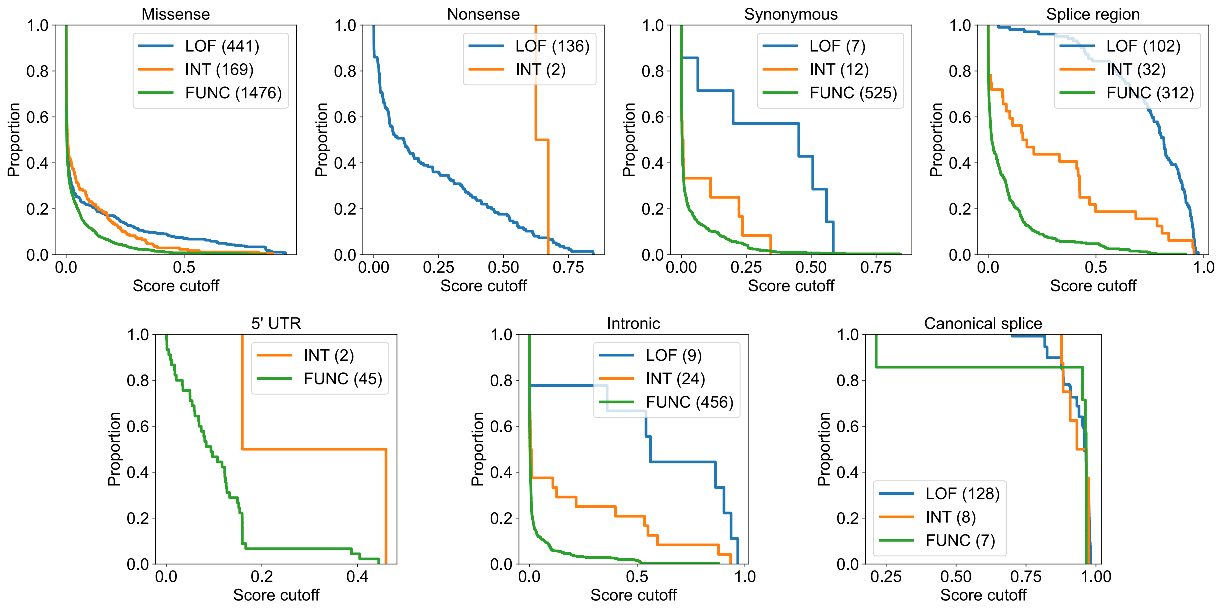


**Supplementary Figure 4. Survival function plots of tested *BRCA1* variants**. Survival function plots of *BRCA1* variants in different annotation bins as a function of predicted splicing effects. The variants are separated by their classification as loss-of-function (LOF, blue), intermediate (INT, orange), or functional (FUNC, green). We observe a huge enrichment of LOF variants among variants with large predicted splicing effects for several annotation classes, with splice regions the most enriched and missense the least enriched. Functional data are from Findlay et al., 2018.

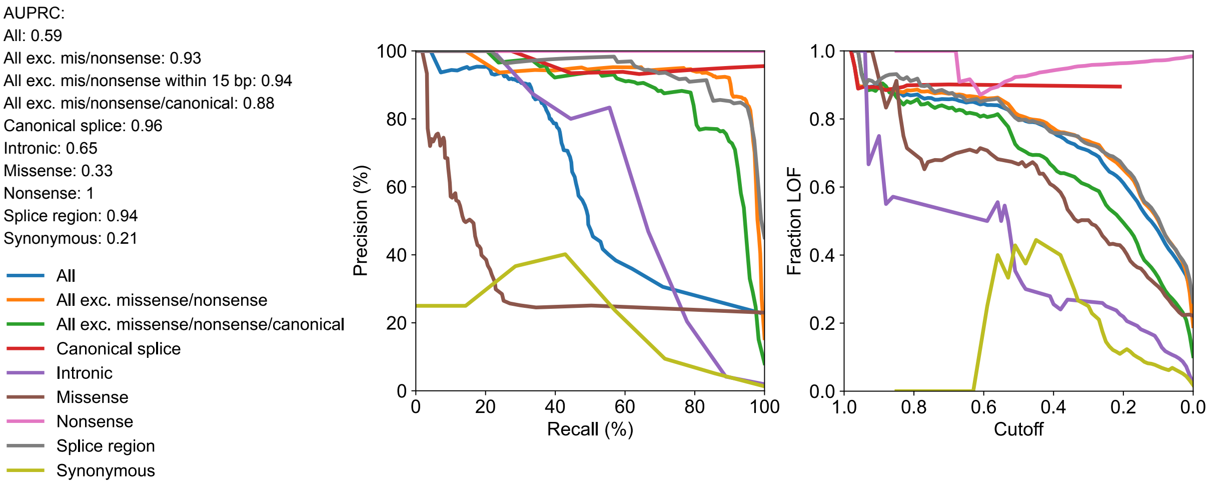


**Supplementary Figure 5.** ***BRCA1*** **predictions**. Left panel: Precision-recall curves, one for the variants in each annotation bin, representing the precision and recall for using Pangolin predictions to distinguish loss-of-function (LOF) variants from functional variants. Right Panel: Line plots showing the fraction of variants classified as LOF as a function of Pangolin's predicted effects on splicing. Despite a poor AUPRC for missense variants, Pangolin maintains a roughly 60% and 80% precision for variants with predicted effects on splicing greater than 0.4 and 0.8 respectively.

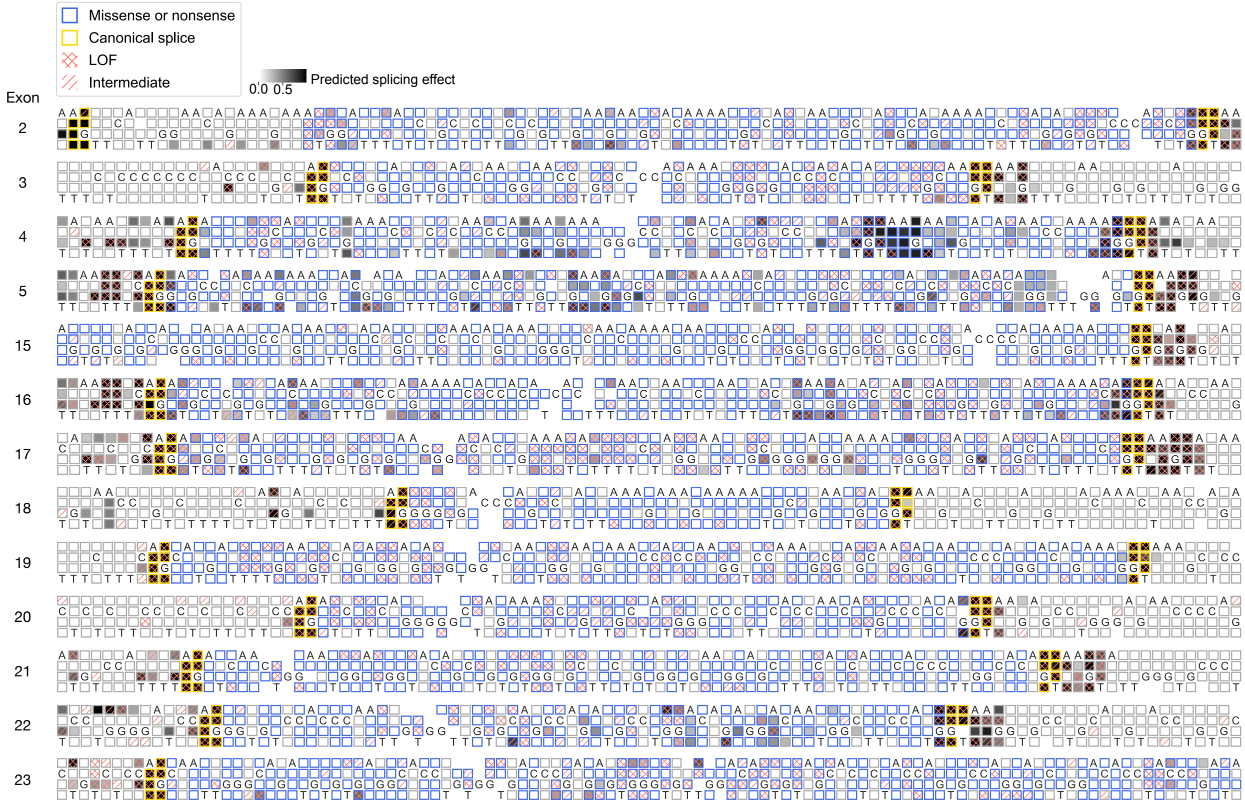


**Supplementary Figure 6.** ***BRCA1*** **predictions**. Predicted splicing effects of *in silico* mutations in or flanking 13 *BRCA1* exons from Findlay et al., 2018. Mutations identified to be LOF or to have intermediate phenotypes, as well as mutations that are missense or nonsense, or affect the canonical splice sites are annotated.

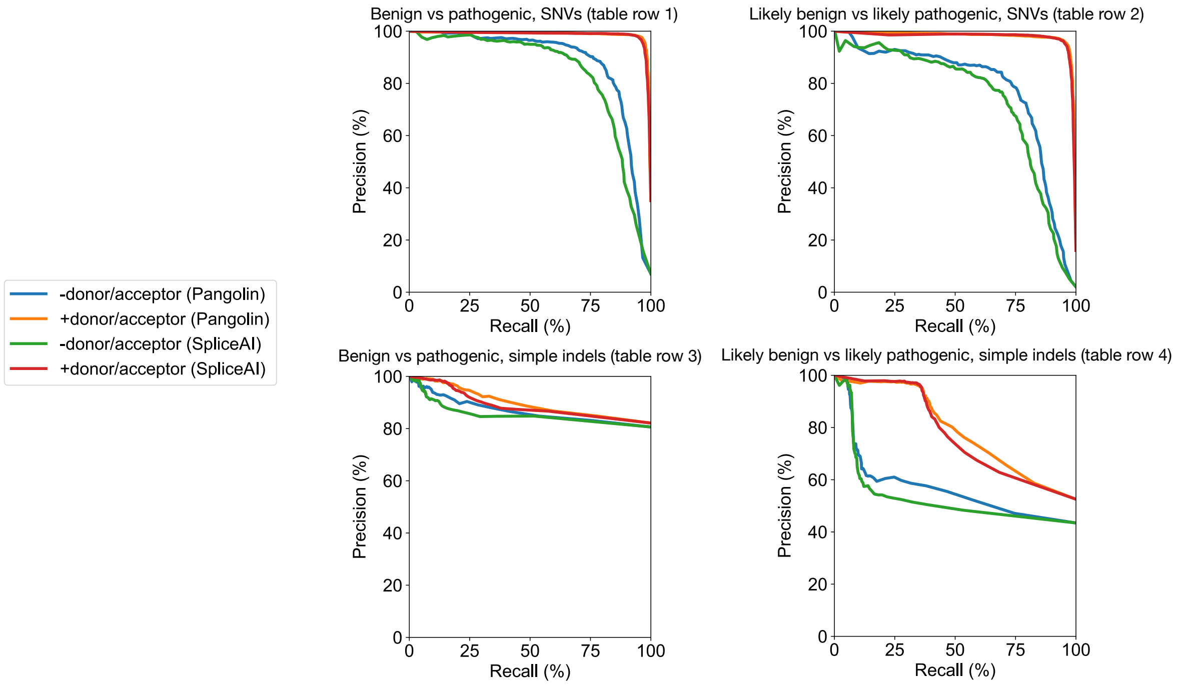


| | | +donor/acceptor | | | | -donor/acceptor | | | |
|---|---|---|---|---|---|---|---|---|---|
| | | AUPRC, Pangolin | AUPRC, SpliceAI | N (likely) pathogenic | N (likely) benign | AUPRC, Pangolin | AUPRC, SpliceAI | N (likely) pathogenic | N (likely) benign |
| SNVs | Benign vs pathogenic | 0.99 | 0.99 | 5,975 | 11,056 | 0.88 | 0.84 | 833 | 10,999 |
| | Likely benign vs likely pathogenic | 0.98 | 0.98 | 5,386 | 28,299 | 0.78 | 0.74 | 590 | 28,209 |
| Simple indels | Benign vs pathogenic | 0.90 | 0.89 | 5,089 | 1,109 | 0.87 | 0.85 | 4,410 | 1,061 |
| | Likely benign vs likely pathogenic | 0.79 | 0.77 | 1,819 | 1,644 | 0.57 | 0.53 | 1,208 | 1,573 |

**Supplementary Figure 7. Precision and recall for pathogenic versus benign variant classification using Pangolin versus SpliceAI**. Top: Precision-recall curves for distinguishing ClinVar variants annotated as pathogenic versus benign (left) or likely pathogenic versus likely benign (right) for SNVs (top) and simple indels (bottom, indels where the reference or alternative allele is only a single base) using Pangolin and SpliceAI, including (+donor/acceptor) or excluding (-donor/acceptor) variants in canonical splice sites. Bottom: Table displaying AUPRC for each precision-recall curve, and the number of variants in each of the classes represented in the curves.

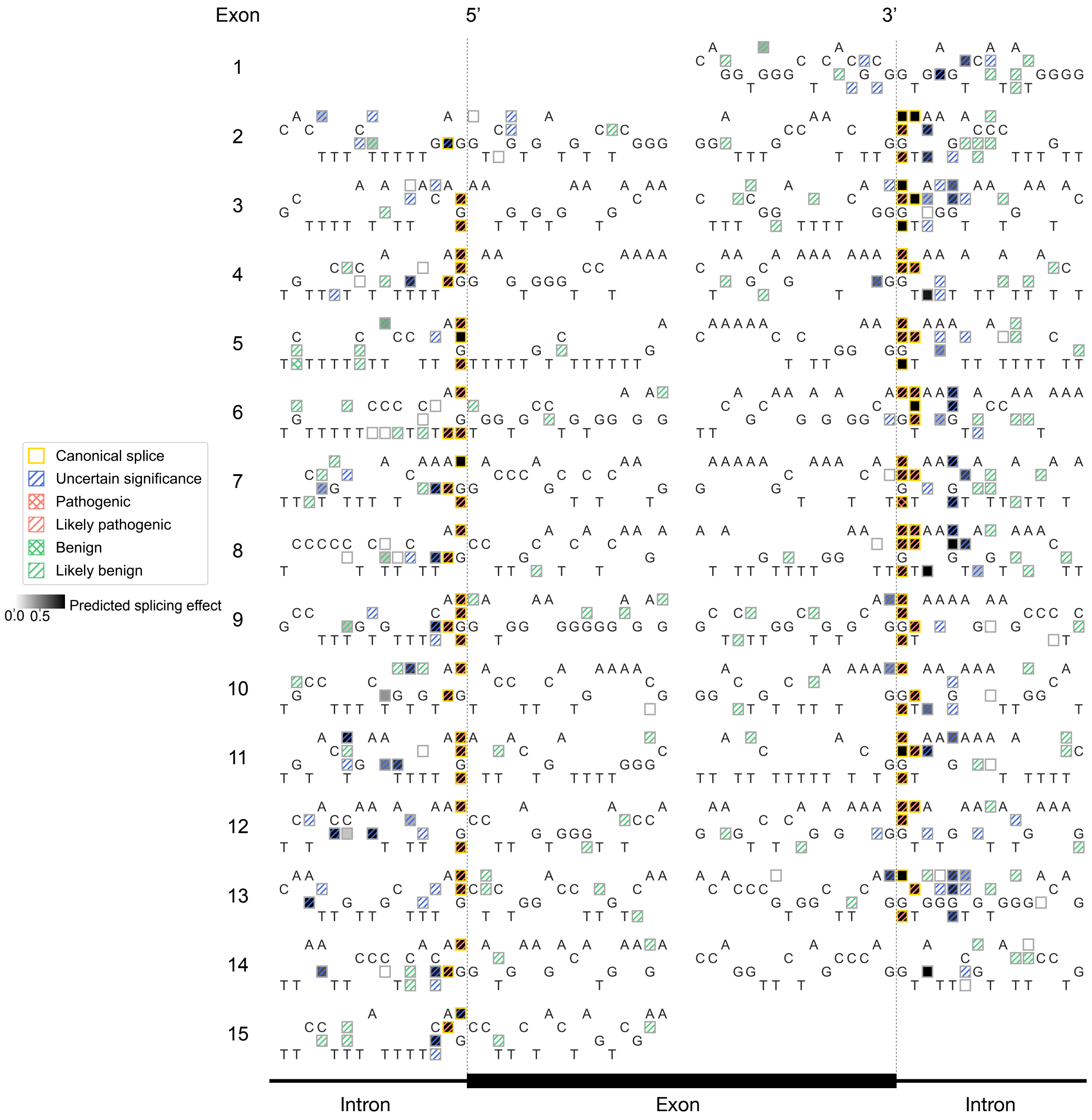


**Supplementary Figure 8. Predicted effects of variants in *CHEK2*.** Many variants of unknown significance, as labeled by ClinVar, are predicted to impact splicing and therefore are likely pathogenic.

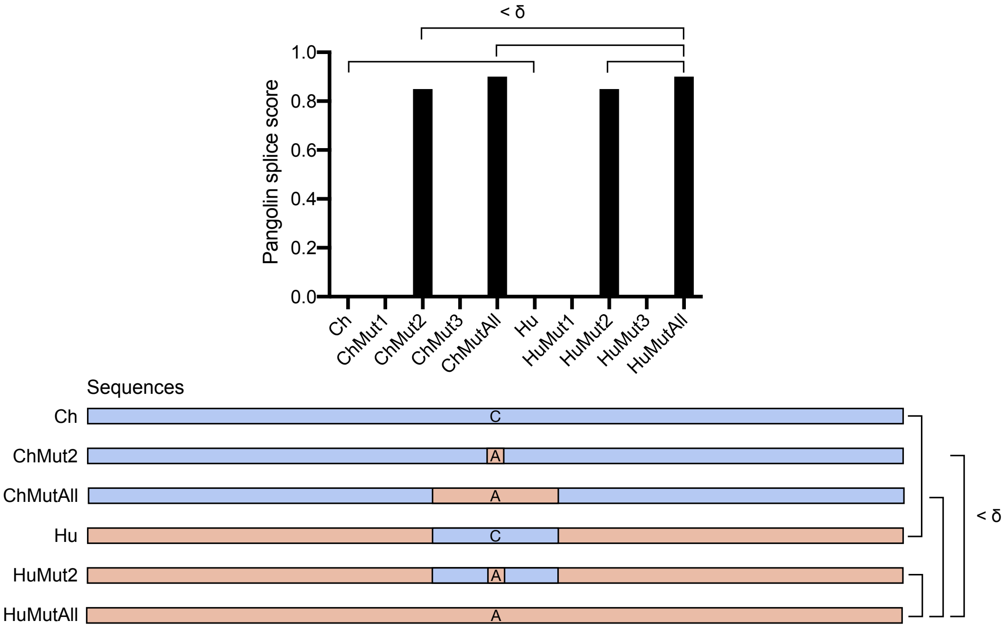


**Supplementary Figure 9. Schematic for identifying single causal variants**. The schematic shows a hypothetical example where three SNVs (Mut1-3) cause in a large increase in the predicted usage of a splice site; specifically, Mut2 is predicted to be the only causal SNV. Ch is the chimp sequence ±5000bp of the splice site, ChMut2 is Ch with Mut2, and ChMutAll is Ch with Mut1-3. Hu is the human sequence ±5000bp of the splice site but with the region near the splice site (±100bp) replaced by the chimp sequence. HuMut2 and HuMutAll are Hu with Mut2 and Mut1-3 respectively. We say that Mut2 is the single causal mutation for this splice site if $|\text{score}_{\text{Ch}} - \text{score}_{\text{Hu}}| < \delta$, $|\text{score}_{\text{ChMutAll}} - \text{score}_{\text{HuMutAll}}| < \delta$, $|\text{score}_{\text{ChMutX}} - \text{score}_{\text{HuMutAll}}| < \delta$, and $|\text{score}_{\text{HuMutX}} - \text{score}_{\text{HuMutAll}}| < \delta$ for X = 2 but not for X = 1 or X = 3.

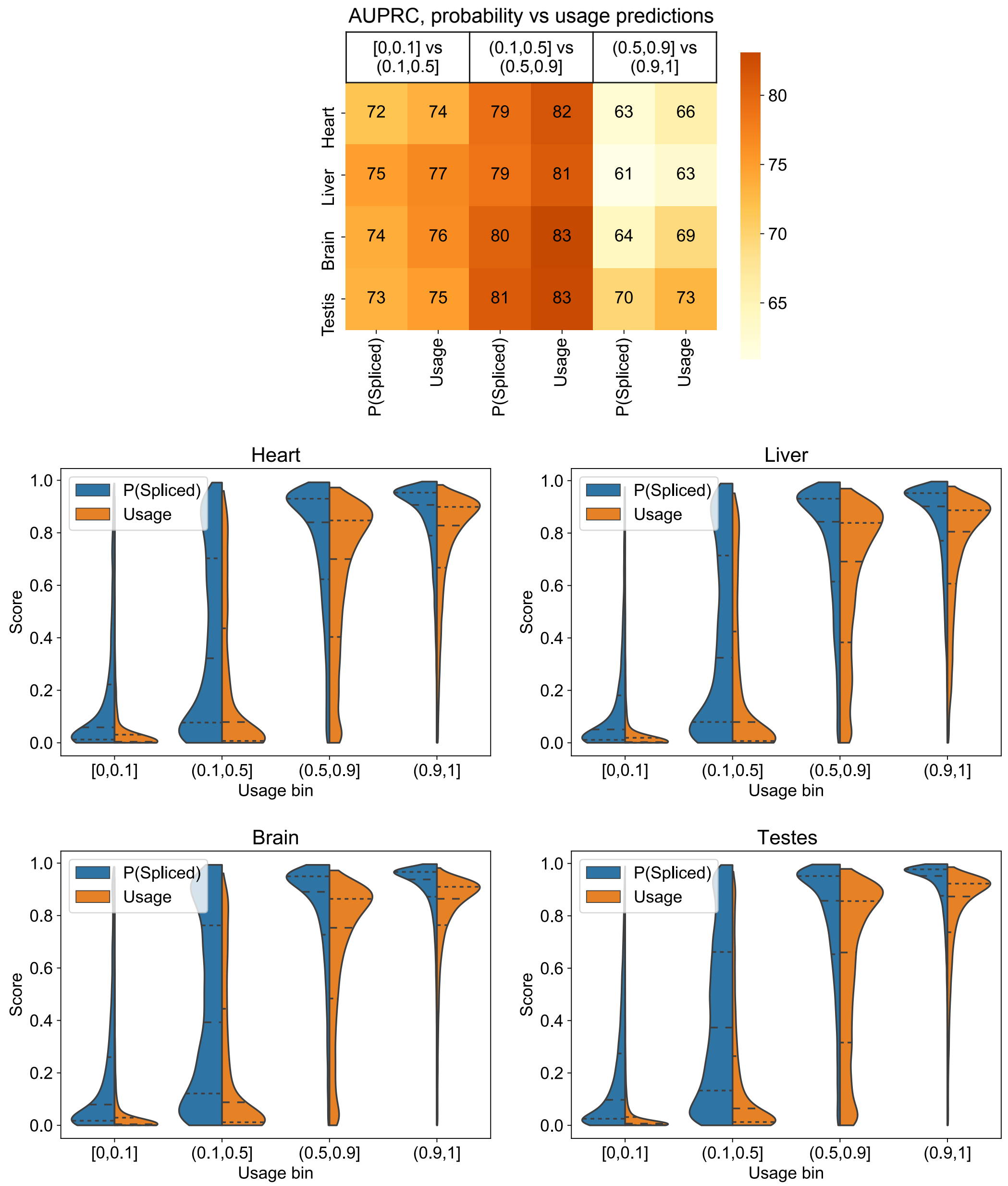


**Supplementary Figure 10. Comparison of training on binary classification of splice sites versus continuous usage estimates**. Heatmap shows AUPRC for classifying splice sites in different usage bins using Pangolin's predictions of the sites' usage and of P(Spliced), the probability that the sites are spliced. As expected, directly training to predict usage improved performance as compared to using the probability that a dinucleotide is a splice site. Violin plots show the distribution of Pangolin's scores (probability and usage predictions) for sites estimated to be used at different ratios empirically in heart, liver, brain, and testis.

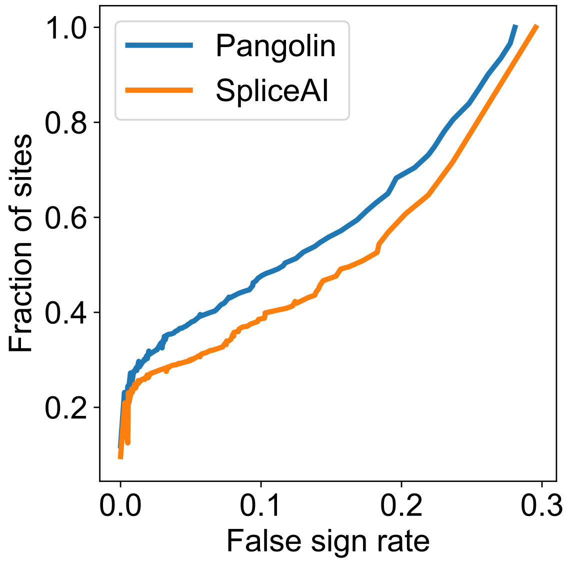


**Supplementary Figure 11. Comparison between Pangolin and SpliceAI for predicting inter-species variation in splice site usage**. Fraction of sites with large differences in splicing ($|\Delta\text{usage}| \geq 0.5$) between chimp and human (1,560 total sites) for which predictions were made at different false sign rates.

|  | All species | Human | All - Human |
| --- | --- | --- | --- |
| Heart (1) | 0.871 | 0.854 | 0.017 |
| Heart (2) | 0.871 | 0.853 | 0.018 |
| Liver (1) | 0.812 | 0.789 | 0.023 |
| Liver (2) | 0.812 | 0.787 | 0.025 |
| Brain (1) | 0.845 | 0.827 | 0.018 |
| Brain (2) | 0.845 | 0.826 | 0.019 |
| Testis (1) | 0.795 | 0.783 | 0.012 |
| Testis (2) | 0.795 | 0.782 | 0.013 |

**Supplementary Table 3. Comparison of AUPRC between models trained on multiple species and on human only**. We compared AUPRC computed over the test set for models trained on human, rhesus macaque, mouse, and rat RNAseq data to models trained on human data only. Models trained on multiple species show consistently better performance.

### References


Baeza-Centurion, P., Miñana, B., Schmiedel, J. M., Valcárcel, J., and Lehner, B., 2019. Combinatorial Genetics Reveals a Scaling Law for the Effects of Mutations on Splicing. *Cell*, **176**(3):549–563.

Cheng, J., Çelik, M. H., Kundaje, A., and Gagneur, J., 2021. MTSplice predicts effects of genetic variants on tissue-specific splicing. *Genome Biol*, **22**(1):94.

Cheng, J., Nguyen, T. Y. D., Cygan, K. J., Çelik, M. H., Fairbrother, W. G., Avsec, Ž., and Gagneur, J., 2019. MMSplice: modular modeling improves the predictions of genetic variant effects on splicing. *Genome Biol*, **20**(1):48.

Findlay, G. M., Daza, R. M., Martin, B., Zhang, M. D., Leith, A. P., Gasperini, M., Janizek, J. D., Huang, X., Starita, L. M., and Shendure, J., *et al.*, 2018. Accurate classification of BRCA1 variants with saturation genome editing. *Nature*, **562**(7726):217–222.
